# Evidence for traveling beta waves in the parkinsonian subthalamic nucleus

**DOI:** 10.64898/2026.07.27.740919

**Authors:** Parsa Nilchian, Alberto Averna, Hayriye Cagnan, Rafal Bogacz, Gerd Tinkhauser, Benoit Duchet

## Abstract

Beta oscillations (13–35 Hz) in the subthalamic nucleus (STN) are a hallmark of Parkinson’s disease (PD), yet little is known about their spatial dynamics. Since traveling waves (spatially propagating oscillations) have been observed in cortical and subcortical structures at different frequencies, we hypothesized that beta oscillations propagate as traveling waves in the STN, likely originating from the motor region of the nucleus. We leveraged multi-contact local field potential recordings from PD patients to estimate pairwise delays between contacts, and asked whether these delays could be accounted for by a simple spherical traveling wave model. Using statistical controls to safeguard against overfitting, we found evidence for spherical traveling beta waves in 11 out of 20 hemispheres, on average during 16% of the recording duration. We further demonstrated that sources are most likely located in the dorsal portion of the motor STN at the group level, and observed a positive correlation between propagation speed and residual motor impairment. Our results suggest that parkinsonian STN beta activity can organise into mesoscopic traveling waves, which has important implications for the pathophysiology of PD and the improvement of sensing-guided DBS.

## 1 Introduction

Beta-band (13–35 Hz) oscillations in the subthalamic nucleus (STN) are a hallmark of the parkinsonian basal ganglia and are closely linked to motor impairment. Neurophysiological recordings from deep brain stimulation (DBS) electrodes revealed pathologically elevated beta activity in the STN of patients with Parkinson’s disease (PD). This elevated beta activity is correlated with the severity of motor symptoms, particularly bradykinesia and rigidity [1, 2, 3, 4, 5]. Moreover, high-frequency DBS of the STN suppresses elevated beta oscillations [6, 7, 8], and the degree of suppression correlates with the improvement in motor symptoms [1, 7]. Yet, despite extensive work on the spectral and temporal properties of STN beta [9, 10, 11, 12, 13, 14] as well as its spectral topography [15, 16], the spatial dynamics of beta within the STN have not been investigated.

Across species and brain structures, rhythmic neural activity often does not manifest as spatially static, but as traveling waves that sweep across neural tissue with a characteristic speed. Such waves have been proposed to support dynamic coordination between neuronal populations and flexible communication among distributed circuits [17, 18, 19]. In the hippocampus, theta waves travel along the septotemporal axis [20, 21]. Beta travelling waves have been linked to the maintenance and use of reward memories in frontal and parietal regions [22]. Traveling waves in the beta band have also been characterised in the human sensorimotor cortex [23, 24], and are carried in the motor cortex by anatomically hardwired circuits between populations of neurons during motor preparation and execution [25, 26]. They are also thought to support the sequencing of proximal-to-distal muscle representation in preparation for reaching in the motor cortex [27]. The spatiotemporal structure of neural oscillations is therefore likely to play a key role in supporting their function.

Whether beta activity in the human STN also forms traveling waves has remained unexplored, largely for technical reasons. In the cortex, high-density microelectrode arrays [20, 22] or electrocorticography grids [23, 17, 28] allow direct visualization of wavefronts as they move across tens to hundreds of recording sites. Conversely, in small deep structures such as the STN, clinical electrodes provide only a limited number of spatial samples that are unlikely to be directly aligned with wave propagation. With such sparse coverage, traveling waves cannot be identified from sequential activation patterns alone, and a model-based approach is needed.

In this work, we hypothesize that STN beta activity propagates as a traveling wave originating from the motor STN. We leverage multi-contact subthalamic LFP recordings and propose a model-based approach informed by delays between contact pairs to study the spatial dynamics of beta oscillations. We find evidence for traveling beta waves, characterise their properties, and provide an initial exploration of their clinical relevance.

## 2 Results

To investigate the spatial dynamics of beta oscillations in the STN, we drew insight from prior observations of traveling waves in other brain structures [21, 29, 30, 28]. Hypothesizing that beta activity in the STN also propagates as a traveling wave, we considered the simplest traveling wave model in three-dimensional space: an isotropic, spherical wavefront propagating at a constant velocity *v* (Fig 1B). In line with the correlation between elevated beta activity and motor symptoms in PD [1, 2, 3, 4, 5], we further hypothesized that these traveling beta waves likely originate from the motor sub-region of the STN. If beta activity in the STN propagates as a spherical wave, it will spread in a predictable manner causing consistent time lags (delays) between contacts’ bursting activity (Fig 1C–D). Conversely, if beta activity is locally generated without coherent propagation, the pattern of delays across contacts should be inconsistent with any simple wave model.

**Figure 1:**
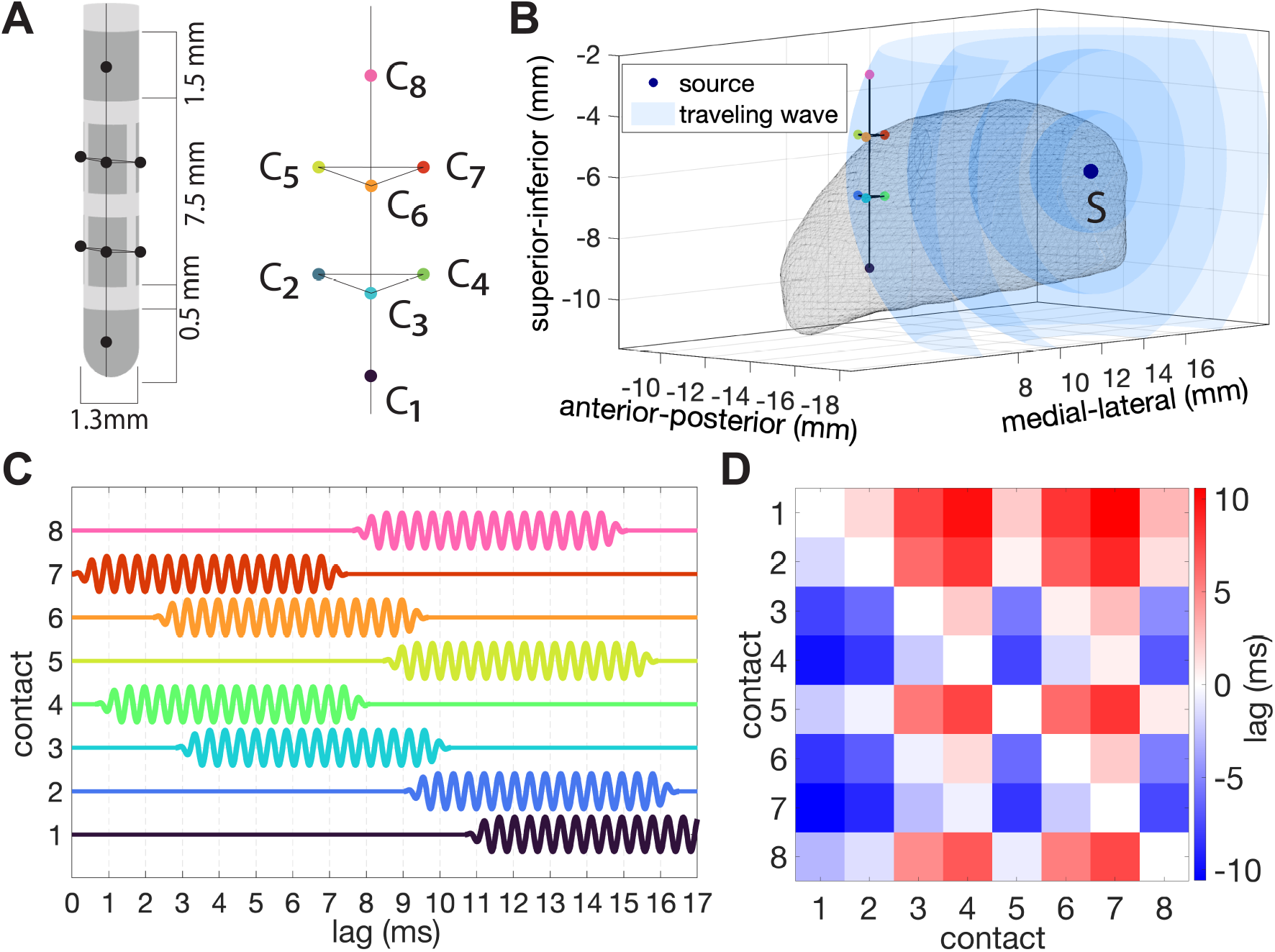
Spherical wave model of STN traveling waves and model of the implanted lead. **A)** Schematic of the implanted VerciseTM Cartesia Directional Lead (Boston Scientific). For simplicity, we used the contacts’ center positions (C*_i_*, *i ∈ {*1*, …,* 8*}*) as the recording locations (colored circles). For directional contacts (C_2_ to C_7_), the center positions are the centers of the contacts’ surface. For omnidirectional contacts (C_1_ and C_8_), the center positions lie inside the lead. **B)** Schematic of the proposed traveling wave model. We hypothesized that STN beta activity may propagate as a spherical wave with a constant velocity originating from a source (S). **C)** Schematic of signals recorded at C_1_ to C_8_ in the presence of a burst of activity propagating from the source S as a spherical wave with constant velocity of 0.1 mm/ms (with contact and source positions as in panel B). The burst of activity will reach C_7_ and C_4_ first. Given that the distances between the contacts’ center positions and the source are fixed, this will result in predictable delays between contact pairs. **D)** Corresponding lag matrix showing the delays between the signals of all contact pairs.

Using synthetic data, we first selected the method best suited to estimate time delays between contacts among a set of candidate methods. We next focused our analysis on LFP recordings from 20 hemispheres across 14 individuals with PD (mean age: 59.0 ± 11.3 years; 10 male, 4 female; see Table A in the supplementary material for participant details). All patients were implanted with a directional, eight-contact (C_1_ to C_8_) DBS lead (Fig 1A). Time delays were quantified in lag matrices (8×8 contacts, Fig 1D) computed from LFPs. We used model fitting to determine whether lag matrices from patient data were consistent with lag matrices expected under the spherical wave model. Having established the consistency of a subset of epochs with the model, we next characterised where these waves originate at the group level, how stable their parameters are, and explored whether these waves may relate to the beta state and motor impairment.

### 2.1 Beta-envelope cross-correlation best estimates STN LFP delays

To assess model validity and reliably estimate the spatial origin and propagation speed of a potential beta wave activity, our approach relies on accurate measurement of time delays between LFP signals. It is however unclear what the optimal method for estimating inter-signal time delays in the beta band is. To identify a suitable method, we therefore systematically evaluated the performance of eight methods (Fig 2A–K) using synthetic data. These methods were based on cross-correlation (XC) or mutual information (MI), and post-processing with varying levels of specificity for beta bursts. While XC is simple and fast to compute, it can only detect (delayed) linear relationships between two signals. Conversely, MI is computationally intensive but was included because it can capture any kind of statistical dependence. We assessed each method using four metrics. First, in synthetic data where ground-truth delays were known, we quantified the absolute error between estimated and ground-truth lags (Fig 2C). Second, we measured the antisymmetry of the resulting lag matrices, which reflects directional specificity of the delays (Fig 2D, J). Third, we assessed the stability of lag estimates across different data epochs, reasoning that a robust metric should yield consistent lag estimates in stationary synthetic data (Fig 2E, K). Finally, we evaluated lag summation consistency error, which quantifies the consistency of time delays across contact triplets based on a (model agnostic) expected additive relationship (Fig 2F, I).

**Figure 2:**
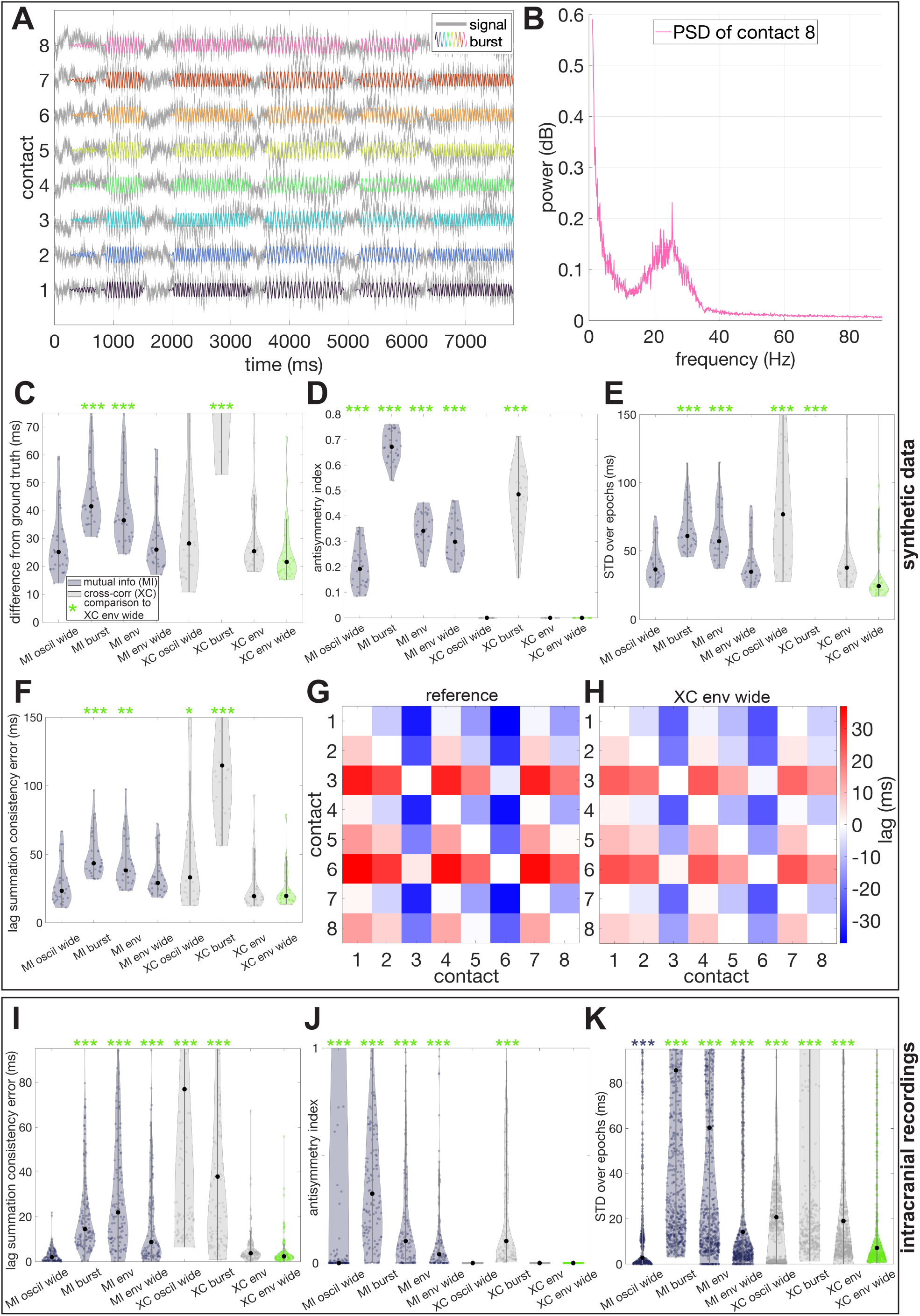
Assessing methods for estimating time delays between STN LFPs. A-H correspond to synthetic LFPs (n = 30), I-K to patient data (n = 20). **A)** Synthetic eight-contact LFPs with known delays. **B)** Exemplary power spectral density of synthetic LFP (contact 8 shown in panel A). **C)** Differences between estimated delays and ground truth time delays in the synthetic data for all methods. **D)** Anti-symmetry index of estimated lag matrices for all methods (range: 0-2, with smaller values indicating greater anti-symmetry; lag matrices are expected to be antisymmetric). **E)** Standard deviation (STD) of estimated time delays computed across 10 s epochs of the data (which should be as low as possible in stationary synthetic data) for all methods. The violin for XC burst is out of range. **F)** Lag sum consistency error for all methods. **G)** Ground-truth lag matrix used in synthetic data (shown for one repeat). **H)** Corresponding lag matrix estimated with XC env wide. **I)** Lag sum consistency error in patient data. **J)** Anti-symmetry index in patient data. **K)** STD over epochs in patient data. In C-F and I-K, *, **, and *** indicate p<0.05, p<0.01, and p<0.001, respectively. Asterisks are green if the method performs worse than XC env wide, black if the method performs better.

To benchmark performance, we first tested all methods on synthetic LFP signals designed to emulate the spectral and temporal characteristics of empirical STN recordings (Fig 2A–H, n = 30 independent repeats). We observed significant performance differences across methods for each of the four metrics (Kruskal–Wallis tests, p < 0.001 for each of Fig 2C-F). Among all methods, XC using the Hilbert envelope of LFPs bandpass filtered *±* 7 Hz around the beta peak (“XC env wide”) demonstrated the best overall performance, ranking highest in three of the four evaluation criteria. Post-hoc Dunn–Šidák comparisons using XC env wide as the reference largely confirmed these findings, although no statistical difference was found with “MI oscil wide” (based on LFPs bandpassed *±* 7 Hz around the beta peak) except for the antisymmetry metric (see Fig 2C-F). Estimated lag matrices closely resembled the ground truth (Fig 2G–H).

We then verified that XC env wide was also adequate in patient data (STN LFP recordings from n = 20 hemispheres), using the three metrics that can still be computed when ground-truth delays are unknown (Fig 2I–K). Consistent with our synthetic data results, XC env wide outperformed the other metrics, producing lag matrices that were more antisymmetric, more internally coherent, and stable across time, with the exception of MI oscil wide (Kruskal–Wallis tests, p < 0.001 in all cases, followed by post-hoc Dunn–Šidák tests see Fig 2I-K). Based on this converging evidence and given the much higher computational cost of MI compared to XC for no significant advantage, we selected XC env wide as the delay estimation method for all subsequent analyses of LFP beta activity.

### 2.2 Evidence for traveling beta waves in the parkinsonian STN

To investigate whether delays between contacts in patient data exhibit the signature of traveling waves, we fitted a spherical wave model to lag matrices from STN LFPs of n = 20 hemispheres of PD patients (obtained using the time delay estimation method identified above focused on the beta band). Hemispheres were included only if at least four contacts exhibited beta activity that exceeded aperiodic noise, and contact pairs were further rejected when showing insufficient pairwise correlation compared to filtered pink noise (see Methods and Table B in the supplementary material). LFPs were segmented into 10-second epochs, a duration chosen to encompass a sufficient number of beta bursts while remaining short enough to assume stationarity. Delay estimation and model fitting were performed separately on each epoch. We then assessed whether lag matrices from patient data were consistent with lag matrices expected under the spherical wave model: an epoch was deemed to show evidence for traveling waves when, under false discovery rate (FDR) control, the data and model lags were significantly correlated with a Pearson’s correlation greater than 0.5, and greater than the 95th percentile of the corresponding surrogate distribution of correlations obtained from fitting to filtered pink noise surrogates (see Methods for more details, and examples in Fig 3A, I).

**Figure 3:**
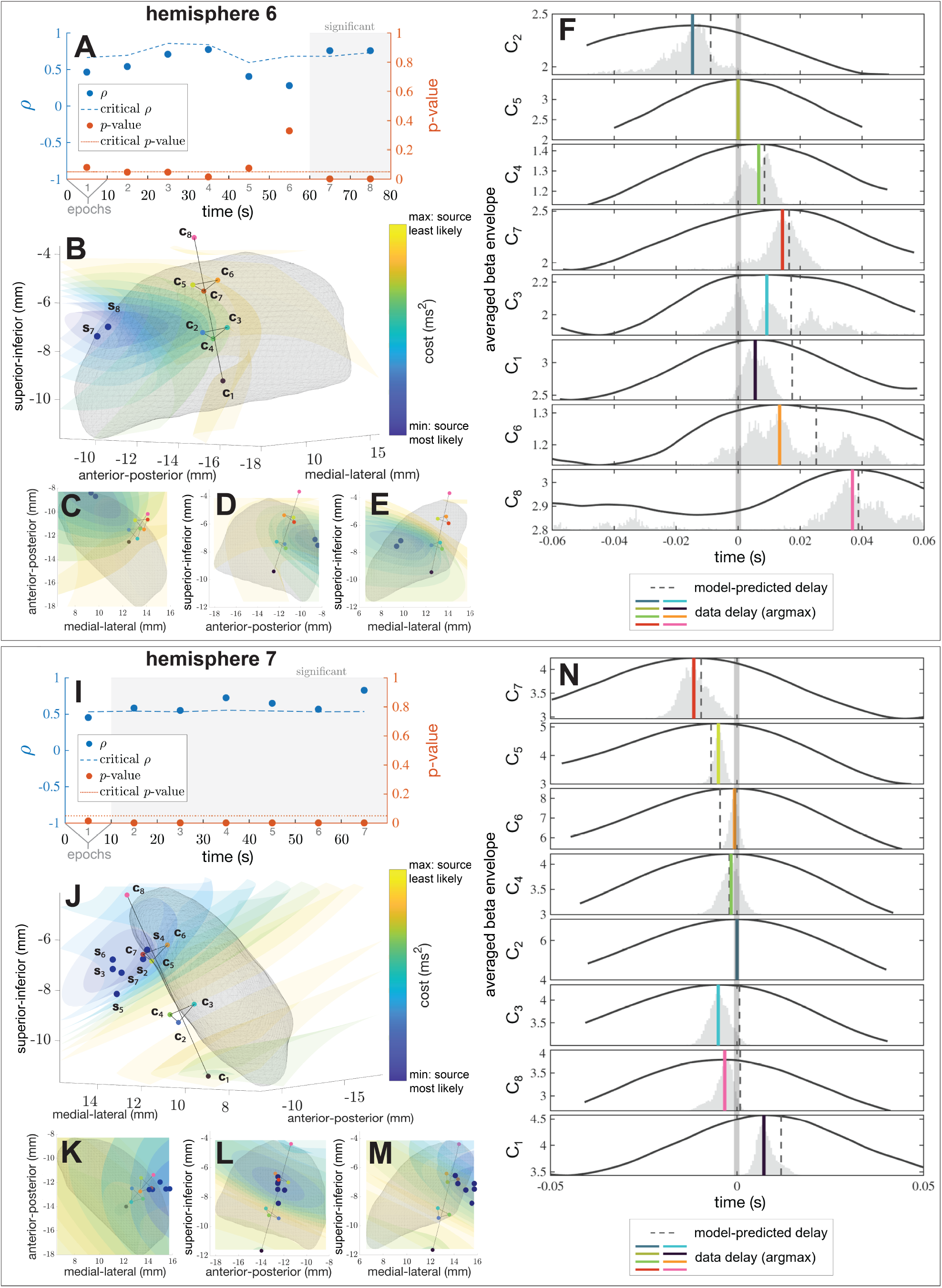
Evidence for traveling beta waves and source localisation in the STNs of PD patients. A, **I)** Overview of significant epochs in hemispheres 6 and 7, respectively. Blue dots represent the Pearson’s correlation coefficient *ρ* between data and model lags, and the dashed blue line indicates the 95th percentile of the corresponding surrogate distribution of correlations obtained from fitting to filtered pink noise surrogates. Red dots show the p-value associated with testing that *ρ* is different from zero, with the 5% significant threshold indicated by a red line. Epochs deemed significant under FDR control are highlighted in grey. **B-E, J-M)** Cost landscape averaged over significant epochs (colored iso-surfaces: areas with lower model cost indicate a better model fit), DBS leads (colored dots and black lines), and estimated source positions (dark blue dots) in hemisphere 6 and 7, respectively. Estimated source positions are shown for all significant epochs (the source corresponding to epoch *n* is denoted S*_n_*). Standardized views are shown in panels C-E and K-M. **F, N)** Visualisation of the wave across contacts for epoch 7 in hemisphere 6 and 7, respectively, obtained by averaging the beta envelope across windows defined based on the beta envelope peaks of a reference contact (C_5_ in F and C_2_ in N). Channels are presented in ascending order of distance to the estimated source. Model predicted delays relative to the reference contact are indicated by dark grey vertical dashed lines, peaks of the averaged data are indicated by colored vertical lines (matching contact color), and probability distributions of the relative timing of the peak of the averaged envelope are approximated using bootstrapped distribution, shown in light grey.

Using this approach, we identified evidence for traveling beta waves in 55% of STNs (11/20 hemispheres). Across all 20 hemispheres, on average 16% of epochs showed evidence for traveling waves (Fig 4A). Two example hemispheres in Fig 3 illustrate the spatial consistency of identified sources across recording epochs. In Hemisphere 6, sources were detected in 2/8 epochs, and their estimated positions were spatially clustered in the dorsal and medial aspect of the STN (Fig 3B–E). In Hemisphere 7, significant sources were found in 6/7 epochs, localized consistently to the lateral-posterior region of the STN (Fig 3J–M). Averaging the beta envelope across windows defined relative to the beta envelope peaks of a reference contact allowed us to visualise the waves identified by the model in these two hemispheres (Fig 3F, N). Having detected traveling waves in a subset of epochs, we next characterised beta wave parameters in these epochs, i.e. the position of the source and the wave speed.

**Figure 4:**
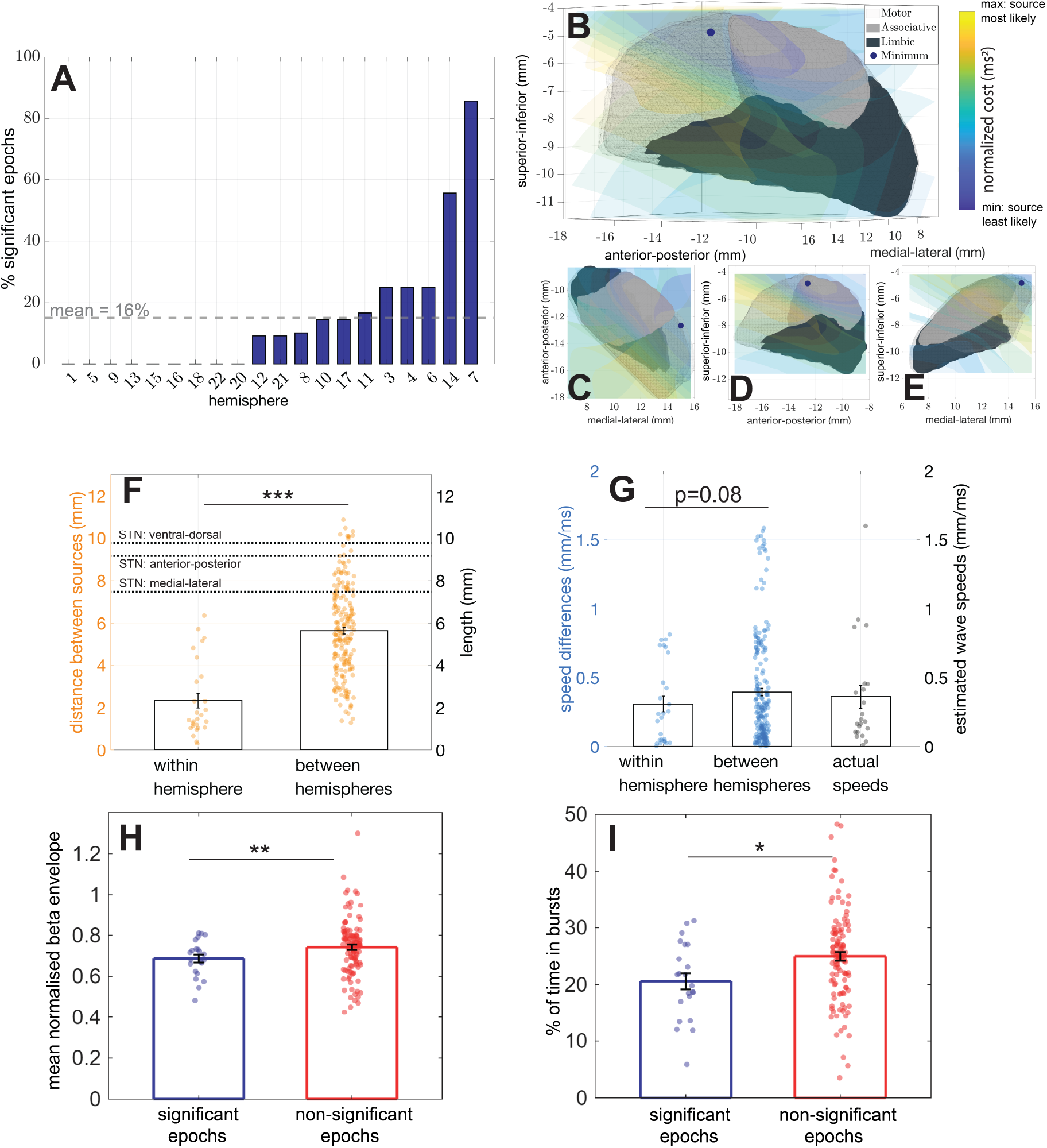
Population-level wave properties. **A)** Overview of the percentage of significant epochs across all 20 included hemispheres. **B-E)** Population-level average cost landscape and STN source position across all significant epochs and hemispheres. The position of minimum cost was in the dorsal motor STN (dark blue dot). Standardized views are shown in panels C-E. **F)** Distance between sources for hemispheres with multiple significant epochs, contrasting the variability within hemispheres and between hemispheres (n within = 27, n between = 204). **G)** Speed differences for hemispheres with multiple significant epochs (n within = 27, n between = 204), and actual speed estimates. **H)** Mean normalised beta envelope during significant (n = 22) and non-significant epochs (n = 113). **I)** Percentage of time spent in beta bursts during significant (n = 22) and non-significant epochs (n = 113). In C-F, *, **, and *** indicate p<0.05, p<0.01, and p<0.001, respectively.

### 2.3 Sources cluster in motor STN while speeds vary

To estimate a population-level origin of beta wave activity, we constructed average cost landscapes for each hemisphere by aggregating the cost maps from significant epochs. These were normalized to the respective maximum cost value per hemisphere and then averaged across hemispheres. In the STN, the resulting group-level cost landscape revealed a consistent minimum in the dorsal motor region (Fig 4B-E), indicating that this subregion may be the most likely initiation site for traveling beta waves.

We next asked whether the estimated source positions and wave speeds were consistent across epochs within individual hemispheres (Fig 4F-G). Among the 11 hemispheres with evidence for traveling waves, four had more than one significant epoch. For each of these, we extracted the estimated source coordinates and propagation speeds across all significant epochs. To assess consistency, we compared within-hemisphere to between-hemisphere variability in both spatial and temporal parameters.

Source positions were highly consistent within individual hemispheres, showing significantly lower variability compared to differences across hemispheres (Fig 4F; p < 0.001). Across epochs within a hemisphere, the average spatial variability in estimated source position was approximately 2 mm—substantially smaller than the overall size of the STN—suggesting stable wave origins across time.

In contrast, estimated wave speeds exhibited greater variability across epochs. Within-hemisphere variability in speed was not significantly lower than the between-hemisphere variability (Fig 4G; p = 0.08), indicating that propagation speed may be more dynamically modulated or sensitive to fluctuations in network state.

Across all 11 hemispheres with significant traveling wave epochs, most estimated propagation speeds fell within a biologically plausible range. The average wave speed was 0.36 mm/ms, well within the range reported in previous studies of traveling waves in both humans and nonhuman primates in other brain structures (0.03–2.4 mm/ms) [30, 25, 23, 28]. These findings support the interpretation that the modeled traveling beta waves may reflect meaningful patterns of spatiotemporal activity in the STN. Therefore, we next explored the relationship between beta waves and the beta state, and whether betawaves relate to motor impairment.

### 2.4 Beta waves, beta state, and motor impairment

We investigated whether epochs with detected travelling beta waves differed in beta burst properties. We found that beta envelope amplitude was significantly higher during nonsignificant epochs (p = 0.006, two-tailed, linear mixed-effects model, n = 135), see Fig 4H. Similarly, the proportion of time spent in beta bursts was higher during non-significant epochs (p = 0.02, two-tailed, linear mixed-effects model, n = 135), see Fig 4I. While these results may indicate stronger beta synchrony when beta activity is not traveling, they are likely impacted by our limited ability to detect traveling beta activity in the STN (see Discussion). Since our limited detection ability would confound the analysis, we do not investigate the relationship between the proportion of epochs with detected waves and motor impairment. However, given the established link between beta synchrony and motor impairment, a faster spread of beta synchrony throughout the STN could be expected to lead to greater motor impairment. We therefore asked whether beta wave velocity was positively correlated with pre-operative movement disorder society unified Parkinson’s disease rating scale (MDSUPDRS) part III scores. Following outlier rejection and averaging across significant epochs and hemispheres within each patient, ON medication UPDRS scores showed a positive correlation with average estimated wave velocity with a trend towards significance (Pearson’s *ρ* = 0.60, p = 0.058, one-tailed, n = 8), but OFF medication scores did not show such trend (Pearson’s *ρ* = 0.44, p = 0.14, one-tailed, n = 8), see Fig S.1 in supplementary material. This was confirmed by linear-mixed effect models predicting the estimated velocity (p = 0.016 for ON medication scores, p = 0.40 for OFF medication scores, one-tailed, n = 21), suggesting that beta wave propagation could play a role in (residual) motor impairment.

## 3 Discussion

### 3.1 Summary of findings

Elevated beta oscillations are a hallmark of PD and are associated with akinetic-rigid symptoms. We investigated the spatial dynamics of beta-band activity in the parkinsonian STN through the lens of traveling waves. We first established in synthetic data that cross-correlating beta envelopes is well-suited to estimate time delays between LFPs from different contacts (Fig 2). Using multi-contact LFP recordings from patients with PD, we then estimated pairwise delays between beta-band envelopes and asked whether these delays could be accounted for by a simple spherical wave model (Fig 1). We found evidence for traveling beta waves in 11/20 STNs, with significant fits in 16% of analysed epochs (Fig 4A). To safeguard against overfitting, we compared model performance to fits obtained from pink-noise surrogates processed identically to the data, and we required model–data correlations to exceed both statistical significance thresholds and the 95th percentile of surrogate fits, under FDR control.

Group-level cost landscapes revealed a consistent minimum in the dorsal motor portion of the nucleus, indicating that traveling beta waves are more likely to originate from this region (Fig 4B-E). Within individual hemispheres that displayed multiple significant epochs, source positions clustered within a radius of roughly 2 mm, substantially smaller than the size of the STN (Fig 4F). In contrast, wave speeds varied more across epochs but stayed within a biologically plausible range. Finally, we found that epochs with detected travelling waves differed in beta burst properties, and that wave speed was positively correlated with motor impairment. Together, these results suggest that STN beta activity can exhibit mesoscopic traveling waves with relatively stable origins and more dynamically modulated propagation speed, which may be associated with motor impairment.

### 3.2 Modeling traveling waves from sparse subcortical recordings

Unlike cortical studies with dense arrays, our deep-brain recordings sample a limited portion of space with an unfavorable geometry, and the underlying wavefront is difficult to observe without prior modelling. To assess whether the observed patterns of delays between contacts were consistent with traveling waves, we therefore considered the simplest traveling wave model with the smallest number of parameters required to describe the data, namely a spherical wave model. This model has only four parameters (Table 1) and is thus less prone to overfitting than more complex alternatives.

**Table 1:** Model parameters and their optimisation bounds. The spherical wave model only has four parameters. Optimisation bounds for wave speed were chosen based on literature describing traveling waves in other brain structures [30, 25, 23, 28, 71, 72]

| Parameter | Symbol | Bounds |
| --- | --- | --- |
| Source position on anterior-posterior plane | $X_s$ | STN A-P: 6.56 to 15.72 mm |
| Source position on medial-lateral plane | $Y_s$ | STN M-L: -18.06 to -8.28 mm |
| Source position on dorsal-ventral plane | $Z_s$ | STN D-V: -11.59 to -4.12 mm |
| beta wave speed | $v$ | 0.01 to 2 mm/ms |

Our spherical model captures the scenario in which the distance between contacts is small compared to the distance from the source, such that the source can be considered pointlike. In reality, the generator of traveling beta waves is likely a spatially extended ensemble of neurons with complex geometry. Our estimated source positions should therefore be interpreted as the centre of such ensembles rather than as single focal points. Despite these simplifications, the inferred wave speeds fall within a biologically plausible range [30, 25, 23, 28] and are in line with the conduction speeds of mono-synaptic unmyelinated connections (0.1-0.6 mm/ms) [31, 32]. In future work with more complex DBS lead design with higher number of contacts (as previously tested [33, 34]), more complicated traveling wave models could be considered to better describe the data (in particular hemispheres where our spherical wave model could not account for the data). For example, a spatially extended source could be approximated by two coherent point-like sources in close proximity.

### 3.3 A mesoscopic traveling beta wave in the STN: implications for parkinsonian pathophysiology

Our findings suggest that, at least some of the time, beta bursts are not spatially static “on/off” events: instead, they emerge at a specific anatomical location (likely in the motor STN) and recruit neighbouring tissue as the wave front advances. In the terminology of a recent theoretical framework [35], these STN traveling beta waves are classified as mesoscopic, type-1 traveling waves. The properties of such waves are thought to be shaped primarily by local connectivity and intrinsic neuronal time constants rather than by long-range interactions alone [35].

Whether these STN betawaves contribute to normal brain function, their potential functional role, and whether/how they are affected by PD pathology are open questions. Unlike theta waves in the hippocampus which are thought to enable phase coding [21], we speculate that beta waves in the STN may orchestrate the activity of STN neurons to maintain the current motor state, in line with the putative role of beta oscillations [36]. Since beta oscillations are hypersynchronised in PD, their spatial propagation as traveling waves could also be exacerbated, which may further reduce the STN’s information coding capacity (proposed to underlie motor symptoms in PD [37, 38]) by coordinating the activity of a large part of the nucleus. Functional ensembles spanning the STN and pallidum have been identified in parkinsonian rats [39] and these waves may even propagate beyond the STN. Given the pathological character of beta hypersynchrony in PD, a faster spread of beta synchrony could be expected to aggravate motor symptoms. Although exploratory, we found evidence that betawave propagation may carry clinical relevance: estimated wave velocity was positively associated with motor impairment in the ON-medication condition, highlighting the potential importance of spatiotemporal features of beta. The STN spatial topography was found to change during motion [16] and across states of consciousness [40]. While comparing the occurrence and properties of traveling beta waves at rest as well as before, during, and after movement may help us understand their possible functional role, comparisons between on and off dopaminergic medication states are likely to bring further insight into the pathophysiology of PD. The spectral topography of higher frequency bands than beta was also found to be predictive of the response to DBS [15], and lower frequencies activity in the limbic region of the STN has been shown to relate to neuropsychiatric symptoms [41]. Whether traveling waves can be identified for these higher/lower frequencies is another open question for future work.

Our results also indicate that beta bursts in the STN may not reflect a single homogeneous regime. While 16% of epochs were well explained by our mesoscopic traveling-wave model, the remaining epochs were classified as non-significant and tended to have higher beta envelope amplitude and a greater percentage of time spent in beta bursts (Fig 4H-I). This pattern is consistent with the possibility that stronger beta synchrony may at times be expressed more globally across the STN in a standing-wave-like manner. However, these results could also simply reflect our limited ability to detect traveling waves with sparse recordings: sequential recruitment of neighbouring tissue may be more easily detectable when beta activity is less frequent and/or weaker – hypersynchrony may mask traveling waves.

### 3.4 Implications for DBS

The development of multi-contact directional DBS leads has enabled site-specific current delivery [42, 43]. However, in current clinical practice physicians have to select the stimulation contact on a trial-and-error basis for each patient, which is very time consuming for medical personnel, fatiguing for patients, and can lead to sub-optimal clinical outcomes. Previous studies have proposed a range of methods to select the optimal stimulation contact. It was shown that beta activity at rest can be used to guide DBS programming [44, 34, 45, 46]. Building on these findings, a combination of spectral LFP features together with statistical learning were put forward as a method to inform contact selection [47]. Other approaches rely on the presence of evoked resonant neural activity [48, 49, 50], spatial filtering [51], or phase-reversal [52, 53, 54, 55] to identify the contact closest to the source of beta activity. In contrast, the source identification method used in this work could enable contact selection by characterising the spatial dynamics of beta oscillations, and localising a source in 3D space in a principled manner. Spatial modeling was also achieved by a model made up of hundreds of thousands of multicompartment STN neurons fitted to patient data [56]. However, the very large computational cost of these complex simulations would be prohibitive for patientspecific clinical applications.

Our amplitude-independent source localisation approach holds distinct advantages over contact selection methods based on amplitude [46, 47]. Stimulation contacts can have slightly different biophysical properties and/or surface sizes and shapes leading to variation in contact impedances, which results in differences in signal amplitude and/or noise level between contacts. This can alter the accuracy of amplitude-based methods. Our method is not affected by this limitation given that it only considers time lags between brain signals recorded from different contacts. Additionally, our method does not require manual selection of contacts within the STN (electrophysiology-based contact rejection only). Head-to-head comparison of our source localisation method with other contact selection approaches will be the subject of future work.

Considering traveling beta wave characteristics may also help to refine spatial parameters beyond current approaches [57, 16]. For example, the size and shape of electric fields could be modified to reflect the traveling wave properties. Accordingly, an electric field designed to suppress a spherical traveling wave would differ from one targeting e.g. planar waves. Due to the progressive nature of PD, plastic changes due to chronic stimulation, changes in medication, or lead movement, we speculate that traveling beta-wave (relative) source locations may be dynamic. Specifically, sources could change shape, shift, or new sources could emerge in addition to already existing ones. Similar source dynamics have already been observed for the starting points of epileptic seizures [58]. Adaptive estimation of the source position on a slow timescale would be feasible given the short duration of data required and the low computation time required (around one minute per hemisphere on a laptop without any optimisation).

### 3.5 Limitations

While our study demonstrates evidence for STN traveling beta waves, it has multiple limitations. First, a greater number of contacts would have improved our ability to detect traveling beta waves – currently likely underestimated – and improved the precision of our source position and velocity estimates. In epochs and hemispheres where we did not detect traveling beta waves, such waves could be present but the geometry of the available recording sites may be ill-suited to constrain the model. While the intermittence of traveling beta waves (detected 16% of the time in this study) could be related to different dynamical regimes across different network states, it is likely to reflect, at least in part, our limited ability to detect them. Second, we considered a spherical wave model with isotropic propagation and a single point source, approximations meant to avoid over-fitting (see above for more details). We also approximated each contact by its centre point (Fig 1), without explicitly modelling its surface area or orientation. Given that estimated source–contact distances are generally large compared to contact dimensions, this is a reasonable first approximation, but incorporating detailed contact geometry could refine source localisation, especially for directional contacts. We also simplified our model by neglecting volume conduction (see justification in Methods), which could have an influence on delay estimation. Third, we did not have access to patient-specific anatomical scans. We were therefore unable to identify source locations relative to patient-specific STNs, and had to consider a generic STN from the DISTAL atlas [59]. Fourth, we only had access to common average reference signals and could not estimate delays between contacts from monopolar signals (for example referenced to the implantable pulse generator). While common average referencing can make the estimation of delays more difficult, we verified using synthetic data that delays between contacts can still be recovered (see Fig S.2 in the supplementary material). Fifth, the surgery-associated “stun-effect” may have affected the spatial dynamics of STN beta, and this study should be repeated with longitudinal chronic recordings to assess stability. Nevertheless, it was shown that contact selection based on intra-operative LFPs may be valid in the long-term [60] and that the spatial distribution of beta power is relatively stable post-surgery [61], suggesting that the impact of the “stun effect” on the spatial characteristics of beta may not be major. Lastly, the association between propagation speed and ON medication UPDRS scores should be confirmed in future studies with larger sample sizes.

### 3.6 Conclusion

By combining multi-contact recordings with a principled traveling wave model, we provide evidence that beta activity in the parkinsonian STN often takes the form of mesoscopic traveling waves likely originating from the motor part of the nucleus. Beyond pointing to the presence of traveling beta waves, our results highlight the potential relevance of spatiotemporal beta features for parkinsonian pathophysiology, suggesting that not only the strength and timing of beta bursts but also their spatial recruitment dynamics may impact motor impairment.

## 4 Materials and Methods

### 4.1 Participants and clinical procedures

#### 4.1.1 Patients

In this study, we had access to data from 27 hemispheres of 17 PD patients (11 male, 6 female) who underwent awake STN DBS surgery at the University Hospital in Bern between September 2017 to September 2019. All patients signed the general consent (local ethics protocol 2017-00551) and data included in this work was used in a previously published study [47]. All patients were implanted with the Boston Vercise Cartesia directional leads (Boston Scientific, Marlborough, MA). At the time of surgery, the patients had a mean age of 58.6 ± 3.1 years and a disease duration of 9.2 ± 1.2 years (for detailed clinical information, including MDS-UPDRS III scores, see Table A in the supplementary material). MDS-UPDRS III scores were obtained during the pre-operative Levodopa challenge. Hemispheres were only included in the final analysis if at least four contacts exhibited beta activity that exceeded aperiodic noise and showed sufficient pairwise correlation (see below), leaving 20 hemispheres from 14 patients as the final study sample.

#### 4.1.2 DBS Surgery and lead description

Dopaminergic medication was withdrawn the night before the surgery (levodopa 12 h before the procedure, dopamine agonists 48 h). To localize the STN, preoperative MRI scans (3T) and CT scans with a Leksell G frame were used in combination with Brainlab iPlan 3.0 Stereotaxy software. The target selection was optimized using microelectrode recordings and selective test stimulation. Leads were implanted bilaterally in all patients. Importantly, the implanted electrodes have a hybrid contact geometry consisting of eight contacts. The bottom (C_1_, bullet tip) and top contacts (C_8_, ring) only allow for omnidirectional recording (Fig 1A). In contrast, contact levels two (C_2_-C_4_) and three (C_5_-C_7_) are segmented, enabling directional recording from three directions angled at 120° from each other. Access to LFP recordings from different STN sites was critical to investigate the presence of traveling waves.

#### 4.1.3 Anatomical contact localization

Leads were reconstructed using the Lead-DBS toolbox version 2.3.2 [62, 63] along with MATLAB 2019b (The MathWorks, Inc., Natick, MA). Using the Statistical Parametric Mapping 12 (SPM12) software and Advanced Normalization Tools [64], preoperative MRI images and postoperative CT images were co-registered and normalized into the Montreal Neurological Institute (MNI) space (MNI 152 NLIN 2009b). To account for error-introducing movements of the brain during lead implantation, coarse and fine masks were used to perform a brainshift correction [65, 62]. Quality of co-registration and normalization were manually checked prior to further processing. Electrode positions and trajectories were reconstructed with semiautomatic PaCER processing and manual corrections [66]. The DiODe algorithm [67] was used to determine lead orientations, which were further checked with post-operative X-ray images. After performing all the above steps, contact locations were described with cartesian XYZ-coordinates in the MNI space, which were then aligned with the anatomical STN from the Distal atlas [68]. All analyses shown in this work were performed in MNI space. To facilitate group analyses, coordinates of contacts implanted into the left STN were converted to right STN analogues with the Lead-DBS toolbox.

### 4.2 Signal processing

#### LFP recordings

Upon placement of the lead in its final position and fixation to the skull, LFPs were recorded during awake DBS surgery from all eight contacts simultaneously. As detailed in the original publication [47], LFPs from 27 out of 34 hemispheres were available for further analyses. The LFPs used for this study were exclusively rest data (mean recording duration across hemispheres 90.7 ± 29.7 s). Using common average referencing, recordings were performed with a TMSi-Porti amplifier (Twente Medical Systems International, Netherlands) at a sampling frequency of 2048 Hz. All further analyses were performed in MATLAB 2021b (The MathWorks, Inc., Natick, MA).

#### Hemisphere-specific peak beta frequencies

For each DBS lead contact in each hemisphere, power spectral density (PSD) was obtained using Welch’s overlapped segment averaging estimator. Line noise was removed with a notch filter centered around 50 Hz. The aperiodic component of the power spectra (Fig S.3 in the supplementary material) was estimated using the FOOOF toolbox [69]. Each contact’s beta power above noise was obtained by subtracting the aperiodic noise estimate from the PSD and applying a Gaussian filter with a standard deviation of 6 Hz on the resulting signal. The frequency corresponding to the highest peak in the beta range (13-35 Hz) was then defined as a contact’s peak beta frequency. Each hemisphere’s peak beta frequency was calculated as the median peak beta frequency across all accepted contacts (see below). Defining a peak beta frequency at the level of each hemisphere was useful since LFPs from all accepted contacts of a given hemisphere had to be filtered around the same frequency in order to estimate delays between contacts.

#### Contact rejection based on beta level

Since we were interested in traveling beta waves, contacts without measurable beta power above aperiodic noise (defined here as a relative peak height of 0.3) were discarded (for details on the relative peak height see Fig S.3 in the supplementary material). This threshold of 0.3 was determined empirically upon visual inspection of a subset of 80 contacts from 10 hemispheres.

#### Data lags between contacts

LFPs were bandpass filtered with a second-order Butterworth filter centered on each hemisphere’s peak beta frequency *±* 7Hz (hemisphere-specific peak beta frequencies given in Table B in the supplementary material). The envelope of the bandpass filtered LFPs was obtained as the modulus of the analytic signal constructed using the Hilbert transform. For each hemisphere and each 10 s epoch, normalized XCs were computed to estimate time lags between contacts’ envelope signals. The XC between the (real-valued) signals *J* and *K* from contacts *C_j_* and *C_k_* was computed as

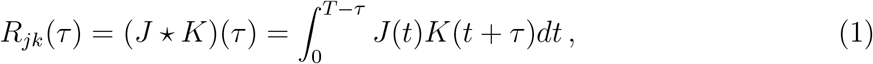

where ★ is the XC operator, *τ* is the time lag, and T is the recording duration. XC coefficients were then normalized according to

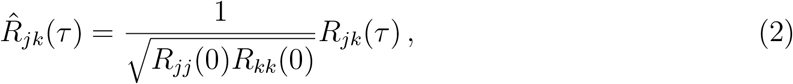

where *R̂_jk_* denotes the normalized XC and *R_jj_*(0) and *R_kk_*(0) are the autocorrelations of signals from contacts *j* and *k* with zero lag. We estimate the time lag in the data 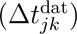 between two contacts (*C_j_, C_k_*) as the delay (*τ*) that resulted in the maximum *R̂_jk_* [70]. Since eight contacts were used in each hemisphere, the resulting lags were summarized in 8 *×* 8 matrices (Fig 1D), with each entry describing the lag between two contacts. By construction, lag matrices are expected to be antisymmetric across their diagonals, which were filled with zeros (Fig 1D).

#### Contact pair rejection based on XC and hemisphere rejection

A second rejection criterion was applied to discard contacts that were not exposed to the beta field e.g. due to their position, orientation (facing away from source), or because other parts of the lead shielded them from the beta-field. For each hemisphere and epoch, we rejected contact pairs with low XC. Specifically, we rejected contact pairs with XC lower than the 95% percentile of the XCs between 50 000 surrogate contact pairs associated with pink noise signals processed in the same way as the data. To prevent overfitting in our models, in addition to the statistical approaches detailed below, epochs with fewer than 20 accepted contact pairs (which is the number of pairs corresponding to five contacts when autocorrelations are ignored) were excluded from the study. The number of five contacts was chosen to have more than half of the contacts available.

### 4.3 Computational modeling of traveling waves

We investigated the hypothesis that STN beta activity may propagate as a traveling wave using the simplest traveling wave model in three dimensions, a spherical wave originating from a point source. If beta activity in the STN propagates as a traveling wave, it will spread in a predictable manner leaving an electrophysiological fingerprint at the contacts (Fig 1B-C). The beta wave will reach contacts closer to the source first, causing a lag (delay) between contacts’ bursting signals.

We neglect the influence of volume conduction (passive spread of electrical potentials through conductive brain tissue). This is justified because volume conduction can be considered to occur instantaneous over the spatial extend of the STN (on the order of nanoseconds), and decays as one over the square of the distance to the dipole considered. Moreover, waves of neural activity are relayed without significant attenuation over distances greater than 10 mm (e.g. in the cortex), thus each contact should feel approximately the same amplitude as the wave sweeps through it.

#### 4.3.1 Spherical wave model

Our spherical wave model was based on the simplifying assumptions that

1. STN travelling waves are spherical,
2. STN travelling waves originate from a single source,
3. and the wave velocity stays constant as the wave spreads.

Our model was constructed with four parameters (see Table 1) describing the traveling beta waves’ source (*X_S_, Y_S_, Z_S_*) and wave speed (*v*). For a given source position (*S*), the Euclidean distance between *S* and a contact (*C_j_*) was obtained according to

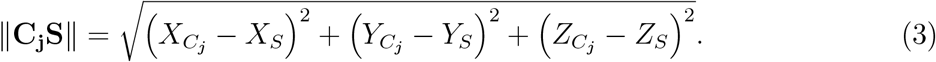

Subsequently, the predicted lag 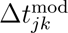 between two contacts *C_j_*and *C_k_* was determined as the difference in travel times from the source

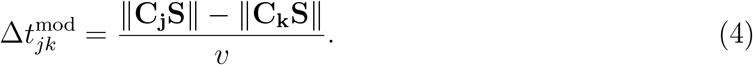

#### 4.3.2 Optimisation procedure

For each hemisphere and each 10 s epoch, the parameters of the spherical wave model were optimised to minimise the discrepancy between time lags predicted by the model and time lags observed in the data. To this end, *N* = 10 local optimizations were performed for each hemisphere and each epoch to increase the likelihood of finding the global optimum. For each parameter, a uniformly distributed random starting point was chosen within set limits (see Table 1). Starting points for wave speed were chosen to be uniformly distributed in log space. Local optimisations were performed with MATLAB’s fmincon function (constrained non-linear optimisation based on interior point optimisation, for algorithmic details see [73]). The parameters resulting in the lowest cost across the *N* optimizations were chosen as the best estimates of the source position and wave speed for the given hemisphere. The cost function was defined as

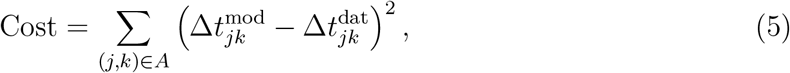

where *A* denotes the set of indices of accepted contact pairs for the hemisphere and epoch of interest.

#### 4.3.3 Cost landscapes

For each hemisphere and each significant epoch, we computed cost across space to map areas of low and high model cost in the STN (areas of low cost correspond to source locations where the spherical wave model fits the data well). The STN was placed at the center of a 15 *×* 15 *×* 15 three-dimensional grid. For each hemisphere, the wave speed was optimized at each individual grid position (*G_i_*) using

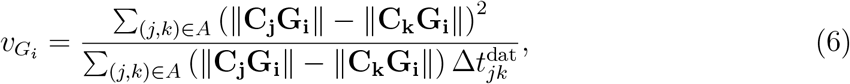

which can be obtained by setting the derivative of the cost with respect to *v* to zero. Iso-cost surfaces were then drawn with MATLAB’s isosurface function.

### 4.4 Model analysis and statistics

#### 4.4.1 Model performance

To provide an intuitive metric for how well our spherical wave model describes the data, we evaluated models using correlation coefficients. The agreement between data and model lags was assessed using Pearson’s correlation coefficient (which we denote by *ρ*). Correlation coefficients closer to one correspond to more accurate models with a lower cost. Since lag matrices (8 *×* 8) are antisymmetric with zeros on the diagonal, only the lower triangle below the diagonals was used for calculating correlation coefficients. For a given hemisphere, an epoch was classified as significant if

1. the spherical wave model significantly fitted the data lags well (*ρ >* 0.5 with *p <* 0.05 for the test that *ρ* is different from zero),
2. the model’s *ρ* was greater than the 95th percentile of the corresponding surrogate distribution of *ρ*’s obtained from fitting to filtered pink noise surrogates (see below),
3. p-values for 1. and 2. survived FDR control at *α* = 5% (applied across all epochs and hemispheres) based on the adaptive linear step-up procedure described in [74].

#### 4.4.2 Pink noise surrogates

A surrogate distribution of correlation coefficients obtained from filtered pink noise was used to control for overfitting. For each hemisphere and each set of accepted contact pairs (to avoid repeating surrogate generation for each epoch), independent realisations of pink noise were filtered as in the data. Similarly to the data, lag matrices were obtained using the XC env wide method, the spherical wave model was fitted to the resulting surrogate lags and the Pearson’s correlation between lags of the fitted model and surrogate data was estimated. This process was repeated 5120 times to construct a distribution of *ρ* values for filtered pink noise.

#### 4.4.3 Averaged wave representation

To obtain an averaged representation of the wave across contacts for a given epoch, we averaged the beta envelope (“env wide”, see below) across windows defined based on the beta envelope peaks of a reference contact, delayed according to model predictions for the contact considered. Envelope peaks were first extracted using a minimum prominence of 0.1 (in units of the envelope normalised by its median during the epoch), and a minimum distance between peaks of 20 ms. To avoid bias, averaging windows were symmetric, centered on the peaks of the reference contact. Their half-width was chosen to include the predicted delayed peak in the contact considered and extended by a further 20 ms. The beta envelope of each contact was averaged across these windows for all detected peaks, yielding an averaged wave representation for that epoch. For each contact (except the reference contact), a boostrap distribution of the envelope argmax position was obtained by resampling peaks within the epoch 5000 times (with replacement). Finally, contacts were sorted based on their distance to the predicted source for visualisation purposes.

#### 4.4.4 Variation in source positions and speeds

To characterise the consistency of source positions across significant epochs within hemispheres, we compared the within-hemisphere variance and the between-hemisphere variance using a pseudo-F statistic given by

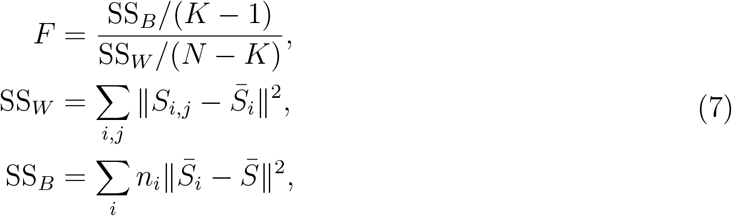

where *n_i_* is the number of significant epochs in hemisphere *i*, *S̄_i_* is the average source position (centroid) across significant epochs in hemisphere *i*, *S̄* is the average source position across all significant epochs and hemispheres, *S_i,j_* is the source corresponding to significant epoch *i* in hemisphere *j*, *N* is total number of significant epochs, and *K* is the number of significant hemispheres. We evaluated statistical significance using 5000 permutations. Source positions were randomly reassigned to hemispheres while maintaining the original epoch count, and the pseudo-F was computed for each permutation. A one-sided p-value corresponding to the alternative hypothesis of greater variance between hemispheres then within hemispheres was obtained from the resulting distribution.

To examine the consistency of source velocity across significant epochs within hemispheres, we also compared the within-hemisphere variance and the between-hemisphere variance. Contrary to source position, velocity is one-dimensional, allowing us to simply perform a one-way ANOVA with epoch velocity as the dependent variable.

#### 4.4.5 Differences between significant and non-significant epochs

We investigated whether epochs with traveling waves differed in beta amplitude and beta burst duration. For each epoch, beta envelope amplitude (“env wide” normalised by its 75th percentile for each channel) was averaged across channels and time. For each epoch, we also averaged across channels the percentage of time spent in beta bursts, defined as the percentage of time the beta envelope was above its 75th percentile (obtained individually for each channel/epoch). We assessed differences between significant and non-significant epochs using linear mixed effect models, with a fixed effect reflecting significance (or lack thereof) of epochs, and a random intercept for each hemisphere. Separate models were fitted for beta envelope amplitude and for percentage time spent in beta bursts.

#### 4.4.6 Correlation between UPDRS scores and estimated wave velocities

To investigate a potential correlation between pre-operative MDS-UPDRS III scores and estimated wave velocities, we considered epochs with significant traveling waves only. We rejected outliers identified by an absolute z-score greater than three, and obtained Pearson’s correlation on the hemisphere- and epoch-averaged data for both ON and OFF medication UPDRS scores. We supplemented this analysis with linear mixed models predicting the estimated velocities with a fixed effect reflecting UPDRS scores (ON or OFF medication) and a random intercept for each hemisphere. Separate models were fitted for ON and OFF medication scores.

### 4.5 Selecting the best method to estimate delays

#### 4.5.1 Candidate methods

To measure the time delay between recordings from different contacts, we considered methods based on normalised XC (see Eqs (1)-(2)), as well as on MI. For methods based on MI, the time delay between two contacts was estimated as the lag that maximises the mutual information between the two corresponding signals (lag search window: −300 to 300 ms, step: 0.49 ms). MI measures how much knowing one variable reduces uncertainty about another (how much information they share). The MI between signals X and Y is defined as

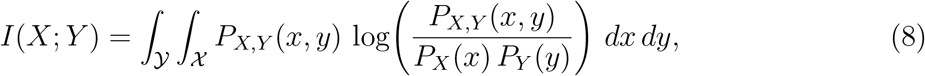

where *P_X,Y_* is the joint probability density function of *X* and *Y* , and *P_X_* and *P_Y_* are the marginal probability density functions of *X* and *Y* , respectively. The MI was computed using [75].

We considered different signals with varying levels of specificity for beta bursts. Signals denoted “oscil wide” were obtained by bandpassing LFPs, with a second-order Butterworth filter centered on each hemisphere’s peak beta frequency *±* 7Hz. Signals denoted “env wide” were obtained as the Hilbert envelope of “oscil wide” signals. Signals denoted “env peak” were obtained as the Hilbert envelope of the bandpassed LFPs, with a second-order Butterworth filter centered on each hemisphere’s peak beta frequency *±* 3Hz. Signals denoted “burst peak” were obtained by applying a contact-specific 75th percentile amplitude threshold to “env peak” signals in order to detect beta bursts [9, 10], and binarising the signal. Periods with amplitudes above the threshold were defined as beta bursts and binarised as ones, the rest of the signal was binarised as zeros.

All eight methods corresponding to the combination of XC or MI and the signals described above were considered for comparison of their ability to estimate time-delays in synthetic data and patient data: “XC oscil wide”, “XC env wide”, “XC env peak”, “XC burst peak”, “MI oscil wide”, “MI env wide”, “MI env peak”, “MI burst peak”.

#### 4.5.2 Metrics

To select the method with the best potential to accurately estimate time delays between contacts, several metrics were employed. The “difference from ground truth” was computed as the sum of absolute differences between the ground truth and estimated lags making up the lag matrix, divided by the number of non-zero entries in the lag matrix. While the difference with ground truth could only be calculated in synthetic data, the following metrics related to expected and desired properties of lag matrices could be computed on both synthetic data and patient data. The “antisymmetry index” was obtained as

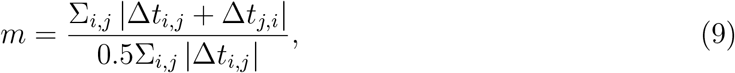

omitting rejected pairs (in patient data). Lag matrices are expected to be antisymmetric, and *m* is 0 if perfectly antisymmetric, 2 if perfectly symmetric. Next, the “STD over epochs” was defined as standard deviation of each Δ*t_i,j_* estimates across epochs (one value per *i, j* per hemisphere/synthetic data run), and omitting rejected pairs in patient data. Finally, the “lag summation consistency error” characterised the departure from the lag sum consistency expected across the lag matrix. Denoting by *t_i_* the time of arrival of an event at contact *i*, and denoting by Δ*t_ij_* the time lag between contacts *i* and *j*, we have (without assuming a particular traveling wave model):

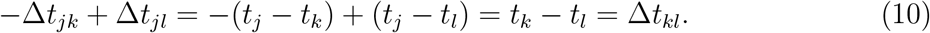

Together with the antisymmetry property, this allows us to construct an expected lag matrix given only one of the rows of the estimated lag matrix. We do this for all the rows, yielding *n_c_* expected lag matrices. For each of the resulting expected lag matrices, we compute the sum of the element-by-element differences between the estimated lag matrix and the expected lag matrix, normalised by the number of elements (we omit rejected pairs when using patient data). We call the normalised sum of these differences across the *n_c_* expected lag matrices “lag sum consistency error”. These four metrics give an indication of how good a method is at estimating pairwise delays, i.e. not being affected by noise and other contributions that might bias estimated delays.

#### 4.5.3 Tests on synthetic data

To evaluate all eight methods using the metrics described above, synthetic data was generated by adding synthetic beta-burst, delayed across contacts according to a given lag matrix, to pink noise (pink noise scale = 100). This process was repeated 30 times using independent lag matrices. Independent lag matrices were obtained by sampling a row of exponentially distributed lags (mean 20 ms), with lags of zero for diagonal elements, and propagating the delays as described above to form a self-consistent lag matrix. Normal noise (mean 0 ms, standard deviation 3 ms) was added to non-diagonal entries of the lag matrix. For each contact, synthetic beta bursts were generated using the Stuart-Landau model (see e.g. [76]) driven by rectangular window inputs. The duration of bursts (window duration) was sampled from a uniform distribution between 50 and 700 ms, and the interval between bursts (interval between windows) was sampled from a uniform distribution between 20 and 1000 ms. Oscillatory activity during bursts was simulated using a Stuart-Landau model with white noise given by

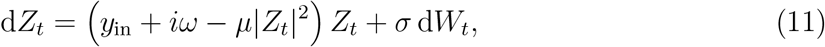

with 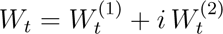 and 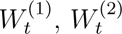 independent real Wiener processes, *µ* = 10, *σ* = 2, and *ω* = 2*πf_β_* where the beta frequency *f_β_* was drawn from a normal distribution of mean 24 Hz and standard deviation 5 Hz (for each burst). The rectangular window input *y*_in_ has amplitude 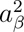 (*a_β_* drawn for each burst from a uniform distribution between 1 and 5), smooth edges based on the logistic function, and added pink noise (scale of 1). For each contact, the burst signal was obtained as the real part of *Z_t_*.

Each repeat was run for 2000 s, sampled at 2048 Hz (as patient data). Methods to estimate delays were applied to 10 s epochs as in patient data. To match the types of signals available in patient data analysis, we subtracted a common average reference from the signal of each contact prior to any data processing. The signal *x_j_* from contact *j* was thus obtained as

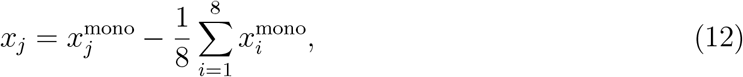

where 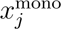 is the monopolar signal simulated as described above. Since beta power was present in all contacts and signals across all contacts were cross-correlated by construction, no contacts/contact pairs were rejected.

#### 4.5.4 Validation on patient data

To validate the choice of method on patient data, all eight methods were applied to patient data. For XC-based methods, method-specific pre-processing was accompanied by the rejection of contact pairs using pink noise surrogates as in the main analysis. For MI-based methods, method-specific pre-processing was accompanied by the rejection of the contact pairs with the worst MI up the the 40th percentile (across all contact pairs and hemispheres).

#### 4.5.5 Statistical analysis

For each metric, values were compared across the eight methods using a one-way Kruskal–Wallis test on ranks, chosen because the distributions were non-Gaussian and heteroscedastic. When the omnibus test indicated significant differences, we conducted Dunn’s post-hoc tests with a Šidák correction for multiple comparisons (multcompare in MATLAB). Post-hoc analyses specifically contrasted each method against the XC env wide method, which served as the reference for assessing delay estimation performance.

## Acknowledgments

We would like to thank Prof. Timothy Behrens for helpful discussions. We would like to acknowledge the use of the University of Oxford Advanced Research Computing (ARC) facility in carrying out this work http://dx.doi.org/10.5281/zenodo.22558. We gratefully acknowledge the contributions of the Bern Parkinson’s and Movement Disorders Centre team, as well as the stereotactic and functional neurosurgery team, for their involvement in patient care.

## Data availability

Anonymized neurophysiological data can be made available upon reasonable request to GT.

