## Supplementary material for "Evidence for traveling beta waves in the parkinsonian subthalamic nucleus"

### A Supplementary tables

| P | H | STN site | Sex | Age (y) | Disease Dur. (y) | Symptoms | MDS-UPDRS III OFF | MDS-UPDRS III ON |
|---|---|---|---|---|---|---|---|---|
| 1 | 1 | L | M | 64 | 16 | akinetic-rigid | 56 | 23 |
| | 2 | R | | | | | | |
| 2 | 3 | L | M | 54 | 11 | akinetic-rigid left | 58 | 27 |
| | 4 | R | | | | | | |
| 3 | 5 | R | F | 70 | 9 | tremor dominant right | 30 | 6 |
| 4 | 6 | L | M | 34 | 7 | akinetic-rigid right | 29 | 10 |
| 5 | 7 | L | M | 55 | 3 | akinetic-rigid right | 40 | 25 |
| | 8 | R | | | | | | |
| 6 | 9 | L | M | 53 | 6 | akinetic-rigid left | 30 | 13 |
| 7 | 10 | L | M | 61 | 9 | akinetic-rigid right | 30 | 6 |
| | 11 | R | | | | | | |
| 8 | 12 | R | F | 53 | 5 | akinetic-rigid right | 47 | 21 |
| 9 | 13 | L | M | 72 | 9 | tremor dominant right | 55 | 14 |
| | 14 | R | | | | | | |
| 10 | 15 | L | F | 60 | 9 | akinetic-rigid, tremor | 41 | 16 |
| | 16 | R | | | | | | |
| 11 | 17 | R | M | 65 | 11 | akinetic-rigid | 38 | 14 |
| 12 | 18 | L | M | 68 | 23 | akinetic-rigid, tremor | 30 | 10 |
| | 19 | R | | | | | | |
| 13 | 20 | L | M | 43 | 11 | akinetic-rigid | 31 | 11 |
| | 21 | R | | | | | | |
| 14 | 22 | R | M | 74 | 12 | akinetic-rigid | 32 | 8 |
| 15 | 23 | L | F | 76 | 4 | tremor dominant | 41 | 27 |
| | 24 | R | | | | | | |
| 16 | 25 | L | F | 33 | 3 | akinetic-rigid, tremor, dystonia | 49 | 8 |
| 17 | 26 | L | F | 62 | 8 | akinetic-rigid | 60 | 27 |
| | 27 | R | | | | | | |
| **Mean ± STD:** | | | | **58.6 ± 3.1** | **9.2 ± 1.2** | | **40.0 ± 2.7** | **19.0 ± 2.6** |

P = Patient, H = Hemisphere, L = Left, R = Right, Dur. = Duration, MDS-UPDRS = Movement Disorder Society Unified Parkinson's Disease Rating Scale. ON/OFF = medication state of patient.

**Table A: Overview of patients' clinical features.** The data presented in this table was adapted from [47].

| H | Rejection 1 | Rejection 2 | Median beta-frequency (Hz) |
|---|---|---|---|
| | Contacts with beta power | <20 contact pairs more correlated than pink noise | |
| 1 | 6 (2) | P | 15.20 |
| 2 | 0 (8) | | |
| 3 | 8 (0) | P | 24.75 |
| 4 | 8 (0) | P | 23.40 |
| 5 | 7 (1) | P | 27.75 |
| 6 | 8 (0) | P | 16.75 |
| 7 | 8 (0) | P | 22.40 |
| 8 | 8 (0) | P | 22.35 |
| 9 | 8 (0) | P | 23.30 |
| 10 | 7 (1) | P | 16.20 |
| 11 | 8 (0) | P | 13.10 |
| 12 | 8 (0) | P | 25.25 |
| 13 | 8 (0) | P | 15.95 |
| 14 | 8 (0) | P | 15.15 |
| 15 | 8 (0) | P | 30.20 |
| 16 | 8 (0) | P | 29.40 |
| 17 | 7 (1) | P | 25.20 |
| 18 | 8 (0) | P | 13.10 |
| 19 | 0 (8) | | |
| 20 | 6 (2) | P | 22.95 |
| 21 | 8 (0) | P | 15.30 |
| 22 | 6 (2) | P | 13.10 |
| 23 | 0 (8) | | |
| 24 | 0 (8) | | |
| 25 | 0 (8) | | |
| 26 | 0 (8) | | |
| 27 | 0 (8) | | |
| **Median \| Mode** | 8 \| 8 | | 20.54 ± 5.71 |

H = hemisphere; P = pass; (N) = number of rejected contacts

**Table B: Overview of rejection steps and median beta frequencies of accepted hemispheres.** Two rejection criteria were applied to select contacts and hemispheres for further analyses. The column corresponding to "Rejection 2" reads pass when at least one epoch was accepted. See Methods for more details.

### B Supplementary figures



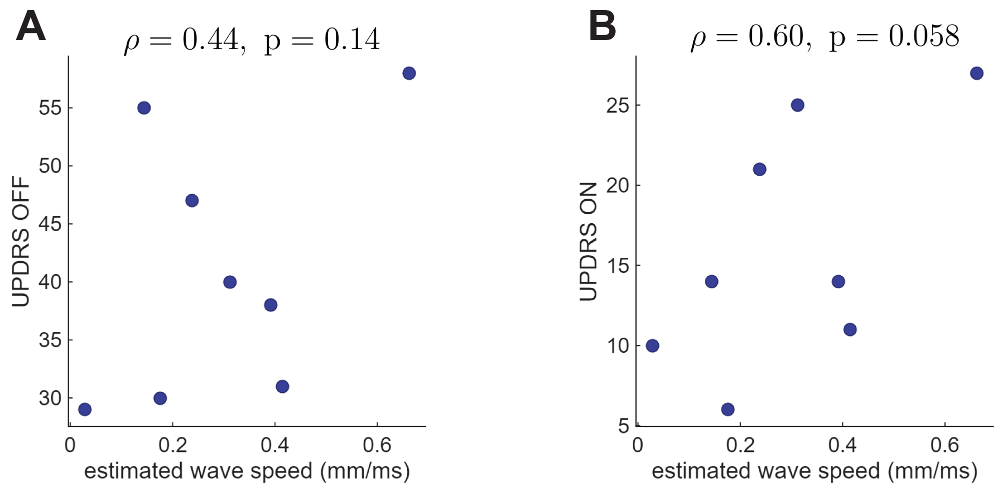


**Figure S.1: Association between estimated wave velocity and pre-operative MDS-UPDRS III scores. A)** OFF medication MDS-UPDRS III score vs estimated wave speed. **B)** ON medication MDS-UPDRS III score vs estimated wave speed. Both panels present data averaged across epochs and hemispheres following outlier rejection.

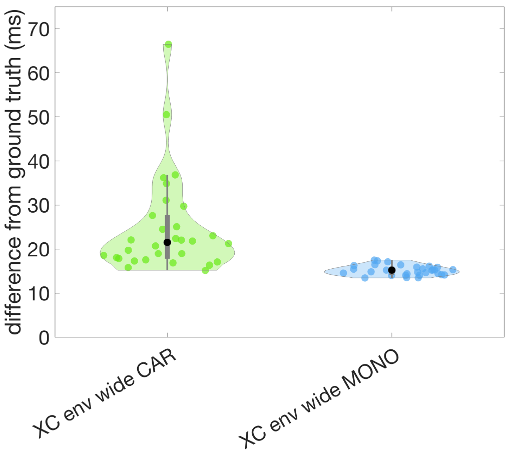


**Figure S.2: Ground-truth error in synthetic data using common average reference signals vs monopolar signals.** Synthetic signals with known delays were generated as in the main text (n = 30). The panel shows the absolute difference between ground truth delays and delays estimated using the XC env wide method, based on monopolar signals (blue, MONO) or signals where a common average was subtracted (green, CAR).

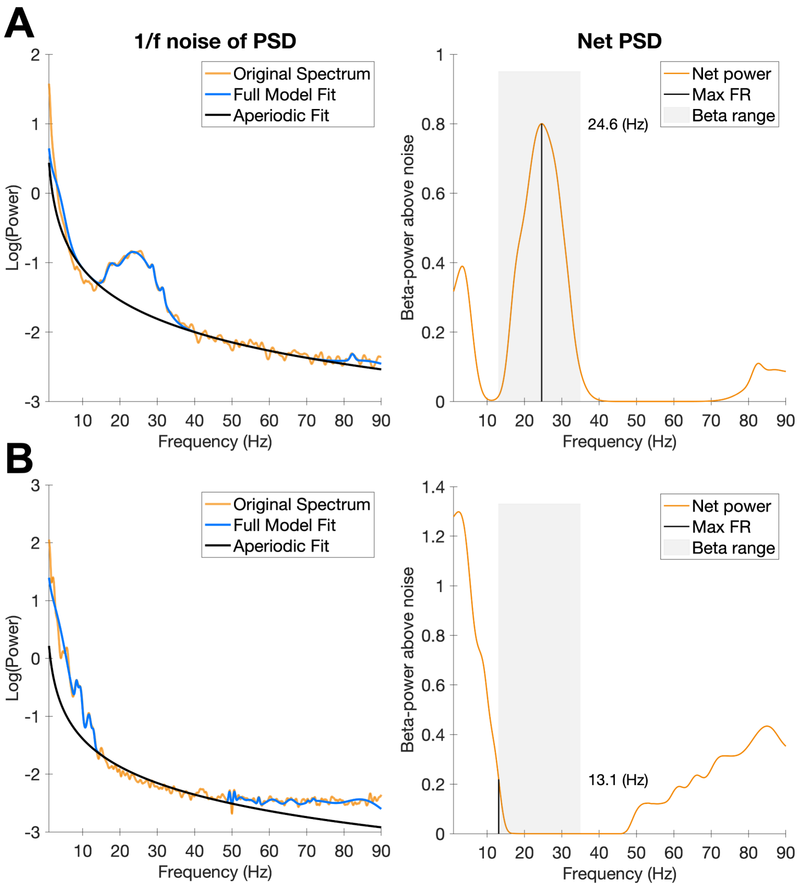


**Figure S.3: Determination of contact-specific peak beta frequencies.** **A)** This contact passed the rejection criterion 1 and was accepted. The PSD in the left panel displays a noticeable peak in the beta range above aperiodic noise levels. Peak beta frequencies were obtained by subtracting the pink noise (black line) from the model fit (blue line) and subsequently extracting the frequency with the highest relative beta power above noise (orange line, right panel). The "model fit" is a smooth representation of the raw data PSD (orange line, left panel). **B)** The PSD (left) and net PSD (right) are illustrative examples of a contact that was rejected according to rejection criterion 1. Noticeably, both PSDs do not display any beta power. H = hemisphere, C = contact.
